# Leveraging remote sensing to distinguish closely related beech species in assisted gene flow scenarios

**DOI:** 10.1101/2024.08.12.607576

**Authors:** Gordana Kaplan, Ariane Mora, Katalin Csilléry, Meredith C Schuman

## Abstract

European beech (*Fagus sylvatica* L.) forests are suffering under increasingly severe and frequent drought. Three closely related, hybridizing beech species, ranging from Bulgaria through Asia Minor and the Caucasus to Iran, offer potential resources for assisted gene flow (AGF) with the aim of increasing the adaptive capacity of European beech forests. However, due to similar morphology and leaf color, as well as hybridization, it is challenging to track the fate of introduced beech genotypes from these related species. Traditional identification methods relying on detailed morphological characterization and genetic testing are labor-intensive and costly, making them impractical for large-scale applications. Using multispectral data from PlanetScope SuperDove, we developed a classification approach that captures phenological differences between the European beech *F. sylvatica* and co-planted Caucasian beech (*Fagus hohenackeriana* Palibin). The approach focuses on key temporal windows and spectral features to optimize classification performance. We evaluated various machine learning algorithms with stratified spatial and temporal cross-validation on data from more than 200 genetically classified individuals in two well-studied sites in France and Switzerland, where Caucasian beech was introduced over a century ago. Our approach was then tested on three different study areas in Germany, where Caucasian beech was also planted, but without specific tree coordinates. Our results reveal consistent temporal and spectral differences during spring and autumn, aligning with budbreak and senescence periods. Most algorithms achieved classification accuracies of 90% and above. The algorithms effectively identified candidate zones for Caucasian beech within or near areas indicated by local foresters. This study demonstrates the potential of high-resolution multispectral satellite imagery and machine learning for scalable classification of closely related and hybridizing species, thereby facilitating forest management in the face of global change.

**Highlights:**

- Remote sensing of unique phenological differences between closely related beech species
- Models with optimized classification parameters differentiated highly similar, introduced trees
- Real-world application to assisted gene flow sites yielded high performance and testable predictions

## 1. Introduction

Climate change and increasing disturbance threaten the integrity of forests (Senf and Seidl, 2022; Patacca, Lindner, Lucas-Borja, Cordonnier, Fidej, Gardiner, Hauf, Jasinevičius, Labonne, Linkevičius et al., 2023). Due to their long generation times, forest trees evolve at a rate that is outpaced by the rate of global change (Aitken and Bemmels, 2016). With the increasing impact of global change on forest ecosystems, there is an urgent need for efficient and scalable methods to monitor and manage tree populations. The planting of species and provenances outside of their existing natural range is considered an adaptive forest management strategy recognized by authorities such as the European Commission. Migration can thus be assisted into new regions that are projected to become suitable for the species niche under climate change (assisted migration, Aitken and Whitlock (2013); McLachlan, Hellmann, and Schwartz (2007)). Alternatively, when a closely related species or a different provenance of the same species is introduced into a population, adaptive introgression can support adaptation in existing forests. Trees harboring different genetic compositions – ideally including heritable traits that are better suited to the future climatic conditions in the target location, referred to as “adaptive provenancing” – can be used for such assisted gene flow (AGF, Aitken and Whitlock (2013); Hällfors, Vaara, Hyvärinen, Oksanen, Schulman, Siipi, and Lehvävirta (2014)). While AGF can be a powerful way to increase forest resilience and adaptive potential, there is also the risk of maladaptation in future generations due to outbreeding depression (Grummer, Booker, Matthey-Doret, Nietlisbach, Thomaz, and Whitlock, 2022). Because the introduced trees are difficult to distinguish visually and may interbreed with the local population, it can be extremely difficult to track the flow of the introduced genetic material. For this reason, foresters have been reluctant to apply AGF at large scales. In contrast, assisted migration of distinct species into formerly uncolonized locations can be tracked more easily, potentially even from space using models to discriminate introduced species and new forest growth from remote sensing data (West, Evangelista, Jarnevich, Young, Stohlgren, Talbert, Talbert, Morisette, and Anderson, 2016), although assisted migration implies a more drastic change to ecosystems.

Climate change is expected to alter the distribution and phenology of many tree species, including the European beech *Fagus sylvatica* L., a keystone forest species in European temperate forest ecosystems (Fang and Lechowicz, 2006; Anderegg, Anderegg, Kerr, and Trugman, 2019; Denk, Grimm, Stögerer, Langer, and Hemleben, 2002). For example, increased temperatures and changing precipitation patterns can stress beech populations, potentially leading to shifts in their distribution and phenology (Wenden, Mariadassou, Chmielewski, and Vitasse, 2020). Eurasian beech species are relevant for studying AGF as an adaptive forest management strategy: European beech forests increasingly suffer from summer drought events that increase both in intensity and frequency (Frei, Gossner, Vitasse, Queloz, Dubach, Gessler, Ginzler, Hagedorn, Meusburger, Moor et al., 2022; Neycken, Wohlgemuth, Frei, Klesse, Baltensweiler, and Lévesque, 2024). Yet it has long been known that eastern Eurasian beeches, long referred to as Oriental beech (*Fagus sylvatica spp. orientalis Lipsky*, Denk (1999a)), exhibit greater morphological (Denk, 1999b) and genetic (Gömöry, Paule, Shvadchak, Popescu, Sulkowska, Hynek, and Longauer, 2003) diversity, which was recently confirmed with modern genetic markers (Kurz, Koelz, Gorges, Carmona, Brang, Vitasse, Kohler, Rezzonico, Smits, Bauhus et al., 2023; Cardoni, Piredda, Denk, Grimm, Papageorgiou, Schulze, Scoppola, Shanjani, Suyama, Tomaru et al., 2024). Accordingly, Eurasian beeches were recently re-classified into four distinct species (Denk, Grimm, Cardoni, Csilléry, Kurz, Schulze, Simeone, and Worth, 2024): the European beech *F. sylvatica*, found from Sicily in the south to southern Scandinavia in the north, and from northeastern Spain in the west to the Carpathian Mountains in the east; the Oriental beech *F. orientalis* Lipsky, found primarily in northwestern Turkey; the Caucasian beech *F. hohenackeriana* Pablin from the Caucasus mountains; and the Hyrcanian beech *F. caspica* Denk & G.W. Grimm from the Hyrcanian forests of Iran and Azerbaijan. Thus, eastern Eurasian beech species are natural candidates for AGF in Europe.

This study exploits a past, non-intentional AGF experiment: one of the newly defined beech species, Caucasian beech *F. hohenackeriana* Pablin (Denk et al., 2024), was introduced into European beech forests in France, Germany and Switzerland the last century (Kurz et al., 2023). Note that Kurz et al. (2023) was published prior to the recent taxonomic revision and thus refers to Caucasian beech as *Fagus sylvatica* subspecies *orientalis* or Oriental beech. While the accurate classification of the two species based on leaf and cupule morphological traits (Denk, 1999b) and/or targeted genetic testing (Kurz et al., 2023) is possible, these methods become labor-intensive, costly, and often impractical at the scale of entire forest stands, multiple stands, or across hybrid zones. Hybridization between the groups in naturally or artificially sympatric regions further complicates their identification, resulting in overlapping leaf morphological traits and phenological stages (Budde, Hötzel, Müller, Samsonidze, Papageorgiou, and Gailing, 2023; Kurz et al., 2023). D’Odorico, Schuman, Kurz, and Csilléry (2023) recently demonstrated the effectiveness of leaf spectroscopy (350-2500 nm) of sunlit top-of-canopy leaves for discerning European from Caucasian beech. Their study, conducted over two summers (2021 and 2022), measured leaf spectral reflectance, morphological and biochemical traits from genotyped adult trees in France and Switzerland. They developed prediction models using partial least squares discriminant analysis (PLS-DA) that distinguished Caucasian from European beech and were most accurate when restricted to either the short-wave infrared (SWIR) region or a suite of traits derived from indices using bands in the visible (VIS), near-infrared (NIR), and SWIR (kappa accuracy of 67-72%). The study highlighted the potential of spectroscopy to scale the discrimination of these very similar species by lessening requirements for resource-intensive genetic analysis and detailed field phenotyping, and providing a basis for upscaling this approach to larger forest areas using remote sensing (D’Odorico et al., 2023).

Through its ability to capture detailed spectral information, remote sensing has revolutionized the study of vegetation and forest dynamics (Willner, Jiménez-Alfaro, Agrillo, Biurrun, Campos, Čarni, Casella, Csiky, Ćušterevska, Didukh et al., 2017). In beech forests, the classification of tree species from distinct genera has recently been demonstrated using satellite remote sensing with machine learning approaches (Grabska-Szwagrzyk, Tiede, Sudmanns, and Kozak, 2024), neural networks (deep learning) on high spatial resolution data (Guo, Li, Jing, and Wang, 2022), or a combination of active and passive data from Sentinel-1 and Sentinel-2 in annual time series combined with forest inventory data (Blickensdörfer, Oehmichen, Pflugmacher, Kleinschmit, and Hostert, 2024). However, the classification of highly similar, interbreeding species groups from remote sensing data remains a largely unsolved problem as current methods still struggle to differentiate groups with similar spectral signatures (Zhong, Dai, Fang, Cao, and Wang, 2024), particularly for species with similar physical characteristics (Li, Hu, Shang, and Li, 2023; Stasinski, White, Nelson, Ree, and Meireles, 2021).

Modern public and commercial satellite imagery provides data with high temporal resolution and increasing spectral and spatial resolution that can be used to detect more subtle differences in the spectral signatures of plant species (Liu, Frey, Munteanu, Denter, and Koch, 2024). Among these options, commercial PlanetScope data, which provides quotas at low or no cost for research purposes, offers several advantages for addressing the challenges such as differentiating between similar tree species with subtle spectral differences and improving classification accuracy in large and complex forested areas. Its high temporal resolution allows for daily revisit times, which is beneficial for capturing dynamic forest phenomena such as spring leaf phenology and autumn leaf senescence, providing detailed temporal datasets essential for phenological studies (Zhao, Diao, Augspurger, and Yang, 2023). In particular, such high temporal density provides more images to choose from within a relevant temporal window, facilitating the selection of high-quality images with low cloud cover. PlanetScope’s high spatial resolution of 3 meters, in comparison to ≥ 10 m resolution of other space-based multispectral platforms, allows pixels to potentially fall on individual tree crowns and cover small forest patches, facilitating more accurate mapping of tree diversity and detecting subtle differences in canopy structure and composition (Reiner, Brandt, Tong, Skole, Kariryaa, Ciais, Davies, Hiernaux, Chave, Mugabowindekwe et al., 2023). However, crown delineation remains a challenge due to canopy overlap and complexity in satellite imagery, compounded by the insufficient 3-meter resolution. Still, the many multispectral bands provided by PlanetScope sensors also allow for relatively detailed spectral analysis in comparison to other space-based multispectral sensors, which is essential for differentiating between species with similar physical characteristics (Kacic and Kuenzer, 2022). A limitation of PlanetScope products and other commercial remote sensing products is that these are not provided with full and public documentation of the methods used in their generation, in contrast to public products which do yet generally have lower spatial, temporal, and spectral resolution (Schuman, Röösli, Mastretta-Yanes, Helfenstein, Vernesi, Selmoni, Millette, Tobón-Niedfeldt, Albergel, Leigh, Hebden, Schaepman, Laikre, and Asrar, 2024). Integrating PlanetScope data with other remote sensing technologies and ground-based observations can significantly improve the performance and reliability of tree species classification and monitoring in diverse forest ecosystems (Wu, Wang, Yan, Song, Chen, Ma, Deng, Wu, Zhao, Guo et al., 2021).

Here, we investigate the potential of high-resolution multispectral satellite remote sensing data to distinguish Caucasian from co-occurring European beech, and develop an approach to classify accurately the two hybridizing species from these data. We aim to (1) understand the spectral differences between the species over time using multispectral bands from satellite-based instruments, (2) select optimal parameters for accurately classifying the species, and (3) develop and evaluate a classification approach that is effective across multiple locations where the species co-occur, starting with European beech stands where Caucasian beech has been introduced for AGF.

## 2. Materials and Methods

### 2.1. Study Areas

We used five study areas, one main study area in France for model development and cross validation and four other sites and Switzerland and Germany for testing (Table 1). We used a beech forest stand located near the village of Allenwiller in northern Alsace, France, at an elevation of 314 meters above sea level (a.s.l., Figure 1A) where Caucasian beech trees were planted in 1923 into a 0.5-hectare plot that is integrated into a larger 4.5-hectare stand of predominantly European beech (Kurz et al., 2023). The species are thus mostly located within proximate but distinct areas, making this an ideal location for conducting spectral analysis and classification studies using remote sensing technologies (D’Odorico et al., 2023). At the time of our investigation from 2019 to 2022, the stand comprised approximately 192 mature trees. Today, all adult trees have been genotyped and their species identity has been determined (Stefanini et al, in prep.). The genotype of a subset of these trees has been published along with a description of the genotyping methods (Kurz et al., 2023; D’Odorico et al., 2023). All species classifications required to reproduce our analysis are provided in the dataset accompanying this paper (see section 7). Adult trees reach heights of up to 40 meters, have a mean diameter at breast height (DBH) of 41 cm, and form a dense, continuous canopy, representing typical mature beech forests in the region. The surrounding area is characterized by a mix of European beech forest and a small spruce plantation to the north.

**Figure 1:**
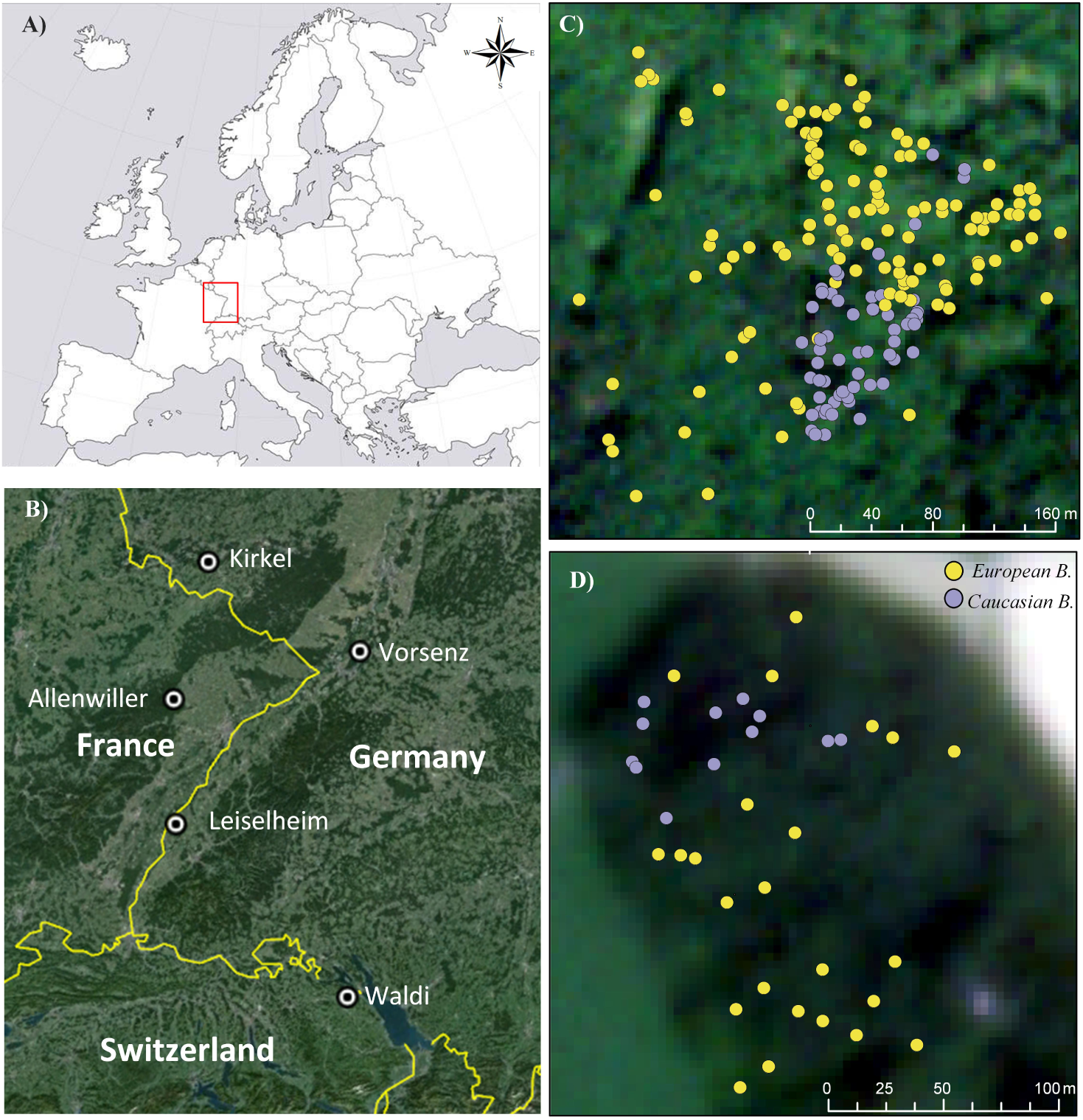
Overview of common beech forest stands used in this study. A) General view of the study areas in the center of the range of European beech (see (Caudullo et al., 2017)). B) Detailed view of the studied sites. Allenwiller is used to study differences between the species and develop classification approaches, whereas Wäldi is used for validation, and Kirkel, Vorsenz and Leiselheim are study areas later used for prediction. Genotyped European and Caucasian beech trees in; C) Allenwiller and D) Wäldi; study areas overlaid on PlanetScope SuperDove satellite imagery (RGB true color combination) from July 2023.

**Table 1.** Central coordinates of the study areas in France (FR), Switzerland (CH), and Germany (D). Coordinates are given using the WGS84 coordinate system.

| Country | Nearby village | Lat (N) | Lon (E) |
| --- | --- | --- | --- |
| FR | Allenwiller | 48°39'00.0" | 7°21'00.0" |
| CH | Wäldi | 47°37'46.0" | 9°06'17.4" |
| D | Leiselheim | 48°07'17.2" | 7°38'30.9" |
| D | Kirkel | 49°17'04.9"N | 7°15'48.7" |
| D | Vorsenz | 49°07'14.3" | 8°27'50.6" |

We conducted the validation on four additional sites: Wäldi in Switzerland, where 36 Caucasian and European beech trees were also previously genetically tested (see Figure 1B), and three beech forest stands in Germany with Caucasian beech planted in a part of the stand, surrounded by European beech. For the latter three sites, we have an approximate coordinate of the stand (Table 1) and a border where the Caucasian beech is expected to be present. Genetic testing has indicated that all of these study sites host Caucasian beech provenances primarily from the Greater Caucasus mountains (Kurz et al., 2023).

### 2.2. Data and analysis

#### 2.2.1. Satellite data and spectral indices

We used PlanetScope SuperDove multispectral satellite images, including coastal blue, red, green, yellow, red, red-edge and NIR spectral bands (see supplementary Table S1). The visible spectrum bands can detect differences in pigmentation, red-edge bands are sensitive to chlorophyll content and canopy structure, and NIR band values are influenced by cellular structure and moisture content over plant canopies (Schlemmer, Gitelson, Schepers, Ferguson, Peng, Shanahan, and Rundquist, 2013; Hoeppner, Skidmore, Darvishzadeh, Heurich, Chang, and Gara, 2020). We also selected spectral indices commonly used for vegetation (see Table 2). Previous work indicated that the combination of spectral band and index values could increase our sensitivity to detect differences between European and Caucasian beech in the VIS and NIR (D’Odorico et al., 2023) and allow us to interpret predictions by relating these to specific physiological and biochemical traits. The band selection for this study was limited to those available for PlanetScope SuperDove, and so we adapted indices by using the closest available bands (Table 2). Nitrogen indices are based on SWIR or red-edge parts of the spectrum (Zheng, Song, Yang, Du, Mei, and Yang, 2022) as such, we included only one nitrogen index which best aligned with the bands available in our dataset. The temporal resolution of our dataset allowed us to observe how differences between the species change with phenology, a distinct advantage of repeat observations from satellite remote sensing technologies (Weisberg, Dilts, Greenberg, Johnson, Pai, Sladek, Kratt, Tyler, and Ready, 2021).

**Table 2.** Spectral indices used in this study.

| Index name | Abbreviation | Formula | Reference |
| --- | --- | --- | --- |
| Nitrogen Index | NI_Tian | $\frac{\text{Red Edge}}{\text{Red Edge} + \text{Blue}}$ | Modified from (Tian, Yao, Yang, Cao, Hannaway, and Zhu (2011)) |
| Normalized Difference Vegetation Index | NDVI | $\frac{\text{NIR} - \text{Red}}{\text{NIR} + \text{Red}}$ | (Tucker (1979)) |
| Simple ratio | SR | $\text{NIR} - \text{Red}$ | (Chang-Hua, Yong-Chao, Xia, Wei-Xing, Yan, and Hannaway (2010)) |
| Triangular Vegetation Index | TVI | $0.5(120(\text{Red Edge} - \text{Green}) - 200(\text{Green} + \text{Red}))$ | (Broge and Leblanc (2001)) |
| Green Index | GI | $\text{Green} - \text{Red}$ | (Smith, Adams, Stephens, and Hick (1995)) |
| Green NDVI | GNDVI | $\frac{\text{NIR} - \text{Green}}{\text{NIR} + \text{Green}}$ | (Gitelson, Kaufman, and Merzlyak (1996)) |
| Photochemical Reflectance Index | PRI | $\frac{\text{Green I} - \text{Green II}}{\text{Green I} + \text{Green II}}$ | (D'Odorico, Schönbeck, Vitali, Meusburger, Schaub, Ginzler, Zweifel, Velasco, Gisler, Gessler et al. (2021)) |
| Optimization of soil-adjusted vegetation indices | OSAVI | $\frac{\text{NIR} - \text{Red}}{\text{NIR} + \text{Red} + 0.16}$ | (Rondeaux, Steven, and Baret (1996)) |
| Transformed Chlorophyll Absorption in Reflectance Index | TCARI | $3(((\text{Red edge} - \text{Red}) - 0.2(\text{Red edge} - \text{Green})) \times (\text{Red edge} - \text{Red}))$ | (Haboudane, Miller, Tremblay, Zarco-Tejada, and Dextraze (2002)) |
| Red edge reflectance parameter I | RERP | $\frac{\text{NIR}}{\text{Green} \times \text{Red edge}}$ | (Datt (1998)) |
| Red edge reflectance parameter II | RERP2 | $\frac{\text{Red}}{\text{Green} \times \text{Red edge}}$ | (Datt (1998)) |
| Simple Ratio RERP | SIRERP | $\frac{\text{Red}}{\text{Red edge}}$ | (Smith, Steven, and Colls (2004)) |
| Normalized Greenness | Norm G | $\frac{\text{Green}}{\text{Red} + \text{Green} + \text{Blue}}$ | (Motohka, Nasahara, Oguma, and Tsuchida (2010)) |
| Leaf Chlorophyll Index | LCI | $\frac{\text{NIR} - \text{Red edge}}{\text{NIR} + \text{Red}}$ | (Le Maire, François, and Dufrene (2004)) |
| Chlorophyll Carotenoids Index | ChCI | $\frac{\text{Green I} - \text{Red}}{\text{Green I} + \text{Red}}$ | (Gamon, Huemmrich, Wong, Ensminger, Garrity, Hollinger, Noormets, and Peñuelas (2016)) |

#### 2.2.2. UAV imagery and tree positions

Tree positions were recorded using a GPS (Trimble Geoexplorer 6000 series GeoXH) and UAV imagery was obtained using a Micasense RedEdge-MX DUAL multispectral camera mounted on a DJI Matrice 210 UAV (DJI Technology Co., Ltd., Shenzhen, China) on July 14^th^, 2022 as described in (D’Odorico et al., 2023). The camera system captures ten spectral bands (see supplementary Table S2), including the visible and near-infrared regions. As the orthophoto had very high spatial resolution (mm), for the purpose of this study, it was resampled to 50 cm spatial resolution (orthophoto available in associated dataset, see section 7).

#### 2.2.3. Spectral Analysis

To discern European from Caucasian beech, we used spectral signature analysis of satellite imagery from April to November, resulting in a dataset across phenological stages with monthly resolution (Shojaeezadeh, Elnashar, and Weber, 2024; Kaplan and Avdan, 2018). The analyses were conducted across four continuous years (2020 - 2023, and additional data set from 2017), and were checked against on-the-ground phenology observations in November 2023. We aimed to select images around the middle of each month but faced frequent cloud cover. As a result, we often could only obtain one image per month. When possible, we included additional images in October and November to better capture the seasonal changes (see supplementary Table S3). Previous research used ground-based spectroscopy measurements with a high number of bands but were limited to measurements performed in July over two years (D’Odorico et al., 2023). We compared our July spectral signature and index results from satellite imagery with field data collected in the same month and year from top-of-canopy leaves (D’Odorico et al., 2023) (Figure S1). To indicate the importance of each wavelength and each spectral index in distinguishing the two species, we used the spectral separability index (SSI) as:

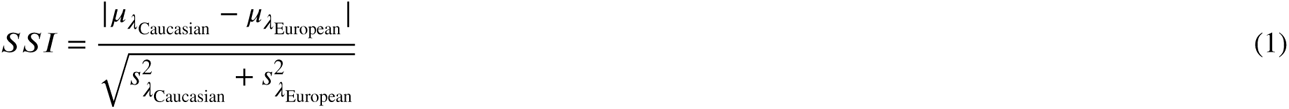

where 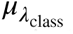 and 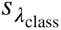 represent the mean and standard deviation of the spectral values for a given bands (*λ*) and class, respectively.

#### 2.2.4. Field observations

We performed a field validation of the phenological differences between the two species. Spring phenology is observed yearly in Wäldi since 2021 (Kurz et al. (2023), Stefanini et al. in prep.), and was recorded in Allenwiller for the first time in spring 2024 (Stefanini et al. in prep.). The autumn phenology was observed in Allenwiller for the first time in 2023 and the results are reported herein.

A team of four observers in two independent groups visited the Allenwiller study site on 11^th^ November 2023. Each observer assessed all trees in the study area. The field observations focused on three main phenological parameters: the percentage of fallen leaves, the color, and the amount of green leaves remaining on the trees. The observers noted the percentage of fallen leaves on each tree. Leaf color was recorded and the amount of green leaves was assessed visually, noting the overall green canopy density. Data from the four observers were averaged to provide a less biased assessment of the phenological state of the species (for the sheet of the field observation, see section 7).

### 2.3. Methods

Our aim was to create a pipeline capable of detecting the subtle differences between European and Caucasian beech using satellite remote sensing imagery. To this end, we evaluated the performance of multiple machine learning (ML) models and tested the effect of temporal resolution on predictions performed using PlanetScope data. Our task was divided into four separate subtasks:

1. Developing an open-source classification pipeline,
2. Identifying optimal features for classification of highly similar species,
3. Training, testing and evaluating ML models for species classification,
4. Using the models to predict each pixel of an image as one of the two species.

The code is publicly available via GitHub, and the data used in the analyses are available at Zenodo (see section 7).

#### 2.3.1. Pipeline and Classification methods

We developed an open-source pipeline for training and testing different machine learning approaches to classify the species based on satellite remote sensing data. Due to the challenges of crown delineation, such as canopy overlap and the insufficient 3-meter resolution, we chose a pixel-based approach and then evaluated classification accuracy given the course-grained data. The pipeline has been developed to meet the specific requirements of our workflow to distinguish European from Caucasian beech using multispectral and multi-temporal satellite imagery. The methodology followed for the classification is given in Figure 2. The pipeline begins with inputting a CSV file containing coordinates and tree classifications. It then aligns these coordinates with the satellite imagery’s coordinate system to ensure accurate spatial matching. The coordinates of the trees were used to identify pixels over either European or Caucasian beech trees, and then extract the pixel values of the spectral data. As several studies have demonstrated improved classification results by incorporating various spectral indices in the classification dataset additional to the raw data(Yaloveha, Hlavcheva, and Podorozhniak, 2021; Estoque and Murayama, 2015), we included the spectral indices given in Table 2. Together with the spectral bands, these features are input into classification models. We evaluated eight classifiers from sklearn: Linear Regression, Nearest Neighbour, Random Forest, Gradient Boosting Machines, Support Vector Machine, MLP classifier, XGBoost, and CatBoost for their effectiveness (Psorakis, Roberts, Ebden, and Sheldon, 2011). Finally, the pipeline outputs the classified data, ready for further analysis or decision-making.

**Figure 2:**
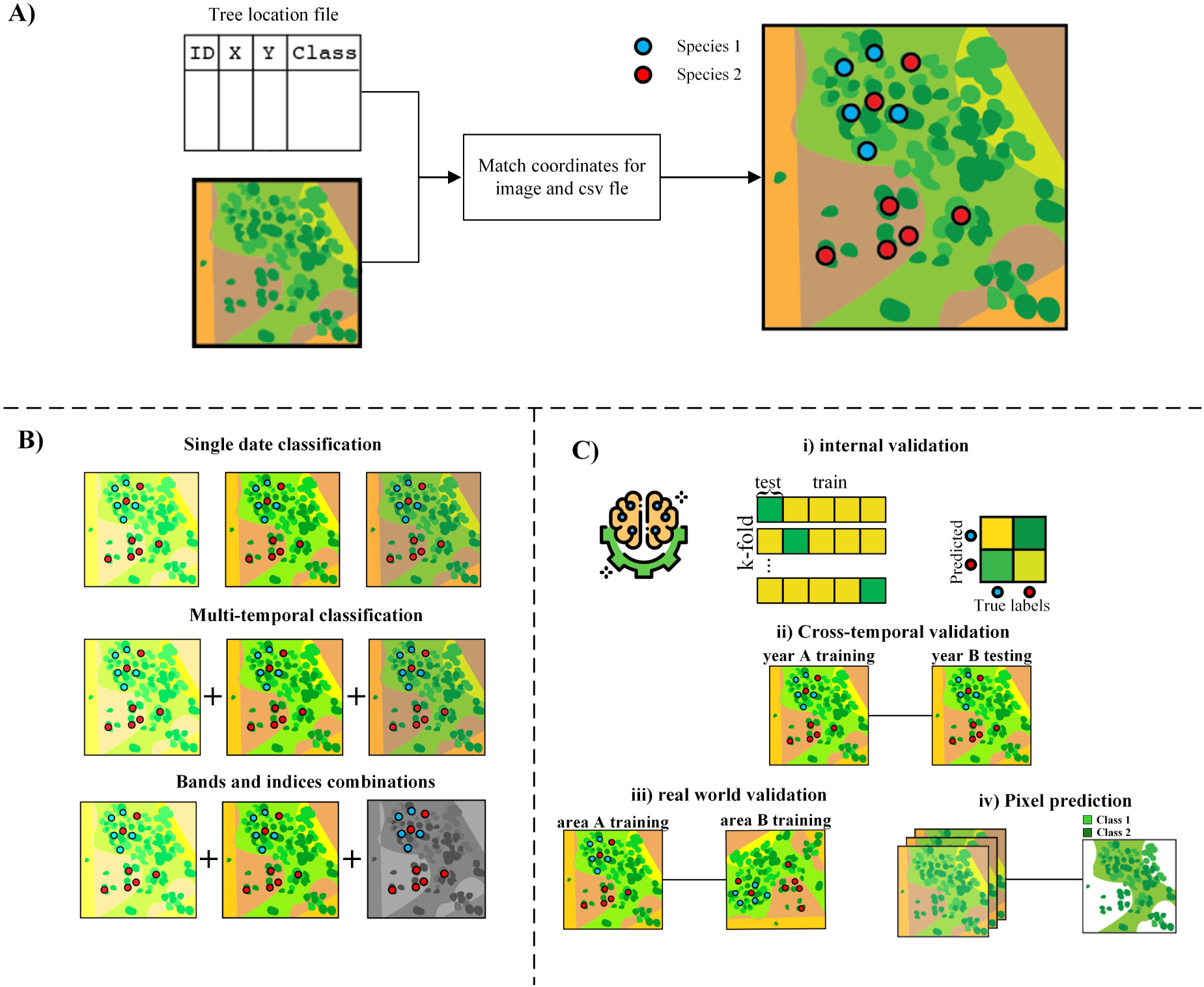
Workflow and methodology implemented for the classification. A) Data section, data preparation. B) Data selection and classification were performed using single-month data, multi-temporal imagery, and a combination of spectral bands and indices. C) Validation of the classification; i) 10-fold internal validation; ii) cross-temporal validation; iii) real-world validation where the Allenwiller dataset has been used as training and Wäldi as testing; iv) Pixel prediction over Wäldi and the three German sites.

We investigated how data from different months influenced the classification performance. With this, we aimed to determine how combining satellite images and spectral indices from different phenological stages affected classification performance. We tested various combinations, including data from a single date, data from multiple dates, and mixed approaches that used both spectral band values and indices from different seasons. Given the limited training data, we only included time points in the final model that reliably distinguished between the species (Van Deventer, Cho, and Mutanga, 2017), to reduce the chance of model over-fitting. Thus, as indicated in Figure 2, we first used single date classification, with each date comprising one set of satellite images from a single day per month from April until November 2023 (eight sets). The classification was conducted with and without the addition of spectral indices. In the second stage, we combined satellite images from different dates based on the results from the single-date classification and the feature selection results. We then selected only the statistically important features and conducted the classification accordingly. This process ensured that models were trained on informative features.

#### 2.3.2. Statistical analysis

To identify optimal features for the species classification, we performed statistical analyses using Python (data and code available in the supplementary files, DATA.txt and t-test-code.txt). We conducted two-tailed t-tests to evaluate the statistical significance of differences in spectral bands and indices between the beech species. To account for multiple tests, we performed Principal Component Analysis (PCA) to assess the dimensionality of the dataset. We then used the number of principal components that explained 90% of the variance to adjust the p-values by multiplying the raw p-value by this number. Features were considered significant if their PCA-adjusted p-values were below a threshold of 0.05.

#### 2.3.3. Cross-validation

To evaluate the classifier performance, we tested multiple cross-validation strategies, including random sub-sampling within single dates, cross-temporal validation across different years, robustness tests with label swapping and dataset balancing, and generalizability assessments across different geographical locations and years. Cross-validation was implemented by randomly partitioning the dataset into training and testing sets for a single date across 10 folds. This repeated random sub-sampling validation ensured that training and testing data were selected differently in each fold (A. Ramezan, A. Warner, and E. Maxwell, 2019). Thus, we could evaluate consistency and reliability over multiple runs, minimizing the bias that might arise from a single data partition. We then tested for generalisability to temporal variation, employing a cross-temporal validation approach, where data from one year served as the training set and data from a subsequent year as the testing set (Meng, Hu, Ren, Zhu, Wang, Liu, and Ma, 2024). By training and testing on data from distinct years, we could evaluate the effect of year-to-year changes which may be due, for example, to plant physiology and phenological variation, on model performance.

Given that the dataset contains unbalanced class labels, we also removed the class bias by sub-sampling the larger class by randomly removing 70 samples (out of 131) from the European beech class, which contained almost twice as many data points. We also evaluated the prediction capacity by swapping labels on pixels so that 50% of European beech pixels were falsely labeled as Caucasian beech pixels, and vice versa. This tested whether our prediction accuracy was actually due to common features within a class, in which case mixing the classes should lead to worse predictions.

To evaluate the generalizability of our approach across different geographical locations, we conducted tests using a dataset of 36 trees from Wäldi, Switzerland that have been genetically classified as European or Caucasian beech. Additionally, we reversed this training and testing setup for exploratory purposes: we used the smaller Wäldi dataset for training and the Allenwiller dataset for testing. This allowed us to explore the capability of our approach when trained on limited data but tested on a broader and more varied dataset. We conducted these testing and training experiments across three different years: 2021, 2022, and 2023. Using this approach, we can infer the generalizability of our model across different geographical and temporal contexts.

#### 2.3.4. Pixel prediction

Having previously focused on pixel classification using specific tree point coordinates from Allenwiller and Wäldi, we trained models to perform pixel predictions on an unlabelled image. For this, we input raster images and predicted the probability of each pixel to be either European or Caucasian beech, facilitating the application of our approach to areas where individual tree coordinates are unavailable. To evaluate this pixel-based prediction approach, we employed the Wäldi dataset but excluded the tree point data and relied solely on the raster images. Afterwards, we incorporated raster data from the three sites in Germany: Leiselheim, Kirkel and Vorsenz, and made pixel predictions of European vs. Caucasian beech.

## 3. Results

### 3.1. Spectral analyses and feature selection

The 8-band PlanetScope satellite data analysis over Allenwiller, France, revealed distinct spectral signatures of European and Caucasian beech over time within each growing season, and the seasonal differences were largely consistent across the investigated years. We investigated 2020 - 2023 and we added four-band PlanetScope data from 2017 for comparison ((see section 7, spectral-values.xsls). We observed that the green, red, and red-edge bands exhibited time-dependent variation, likely related to the seasonal turnover of leaf pigments and overall phenology. European and Caucasian beech were most consistently distinguished by the Green I and NIR bands, with differences becoming larger later in the autumn season (November) (Figure 3) (for all years, see section 7, spectral-values.xlsx). The results for July were consistent with differences in ground-based spectroradiometer measurements (Figure S1) (D’Odorico et al., 2023), except for some discrepancies in the NIR that may be explained by the influence of canopy structure on satellite observations, but not single-leaf observations.

**Figure 3:**
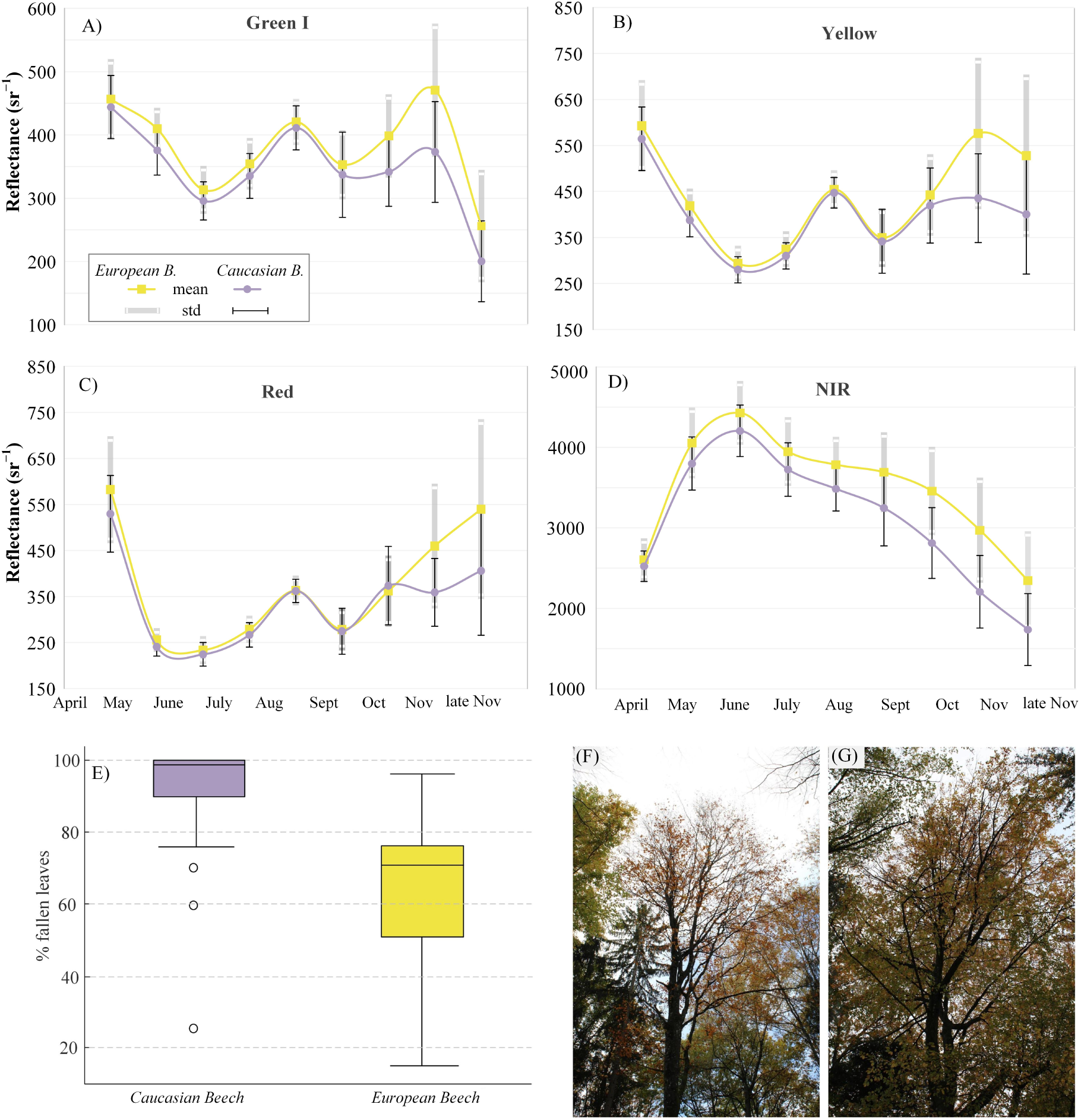
Phenology and differentiation of European from Caucasian beech in PlanetScope superdove imagery. Reflectance values over time for pixels containing crowns of European beech (yellow) or Caucasian beech (purple) from PlanetScope SuperDove images (one image per month), mean and standard deviation, data shown are for 2022, n = 187 pixels (126 for European and 61 for Caucasian Beech) in the (A) Green I; (B) Yellow; (C) Red; (D) NIR bands; and field observation differences between the two species: (E) boxplots showing the percentage of fallen leaves per tree, (F) example photo of an Caucasian beech, and (G) example photo of a European beech.

The statistical analyses supported these observations and showed significant differences in the Green I (t-test, n = 187, t=-3.37, PCA-corrected p = 0.01) and Red Edge (t-test, n = 187, t=-3.19, PCA-corrected p = 0.04). The NIR band differed significantly between pixels over European and Caucasian beech in all of the months except July and August (see section 7, t-test-pca-results.csv). In May and November, there were four and six significantly different bands, respectively, whereas there were two significantly different spectral bands in the other months, suggesting that May and November captured more pronounced differences in spectral characteristics between the species (see section 7, t-test-pca-results.csv).

In the autumn, and especially in November, pixels over European beech canopies exhibited significantly higher reflectance values across all bands, except for the coastal blue, where both species showed similar patterns (t-test, n = 187, PCA-corrected p < 0.000 for bands 2-8; see section 7, t-test-pca-results.csv). Accordingly, field observations in November 2023 for 45 Caucasian and 37 European Beech trees at Allenwiller indicated a clear difference in leaf abscission rates: over 90% for Caucasian beech compared to 50-70% for European beech (Figure 3A). This indicates that the differences observed in late fall between the species resulted from earlier leaf abscission by Caucasian beech. The analysis of the mean temporal values of the spectral indices (see Table 2) showed that for most of the indices, pixels over European beech canopies had higher values for most of the months, namely for NITIAN, NDVI, SR, GI, TCARI, RERP, and RERP2. However, some of these indices showed significant differences in specific months. For instance, NITIAN and RERP2 showed significantly higher values over European beech in November, GI in early November and lower in late November, TCARI in early November, and NORMG and ChCl showed higher values for European beech in October and lower in late November (see Figure S2 and section 7, indices.xlsx). Finally, the SSI analysis indicated the highest separability in autumn, with values ranging from 40 to 75% (Figure S3).

The initial spectral analysis indicated that not all features or time points were equally informative. To select informative features to continue with machine learning, we used the PCA-corrected p-values. The PCA showed that 21 components explained 90% of the variance. Thus, from 207 features (8 band values per month totaling 72 spectral band values, and 15 spectral index values per month totaling 135 index values), 76 of these were found to be significant with PCA-corrected p-values lower than 0.05 (correcting based on 21 independent tests; see section 2.3.2). Of the significant features, 23 were spectral band values, with the largest number in May and November, and the others were spectral index values. The majority of the important spectral index values were from the autumn months, followed by April, where eight (NITIAN, SR, NDVI, GNDVI, PRI, OSAVI, TCARI, LCI) out of fifteen spectral indices were found to be informative in discriminating the two species. These indices provide insight into the phenological differences between the two species, suggesting that in April, European and Caucasian beech exhibit distinguishable differences in their nitrogen content, overall vegetation health, chlorophyll concentration, photosynthetic efficiency, and response to environmental conditions. On the other hand, none of the indices in June and July differed significantly between the species (see section 7, t-test-pca-results.csv).

### 3.2. Classification

#### 3.2.1. Single date classification

The results of single-date classification varied significantly across the year (2023), with April and November showing the best performance. The analysis containing only the spectral band values showed that in April, the average F1 score (harmonic mean of precision: 84% of positive scores which are true positives, and recall: 90 true positive rate) was 0.86, with a range of 0.74 to 0.93, while November had higher F1 scores, averaging 0.88, with a range from 0.81 to 0.93. These months showed a consistent and strong ability to distinguish the species (Figure 4A and Figure S4; see section 7, results-single-date.xlsx and results-single-date-indices.xlsx). In contrast, the mean F1 score in other months was generally lower than 0.85. July, in particular, had the lowest performance, with F1 score values ranging from 0.64 to 0.87 (Figure S5). The indices slightly improved the results of all months, except for April and, in most cases, reduced the range of the scores.

**Figure 4:**
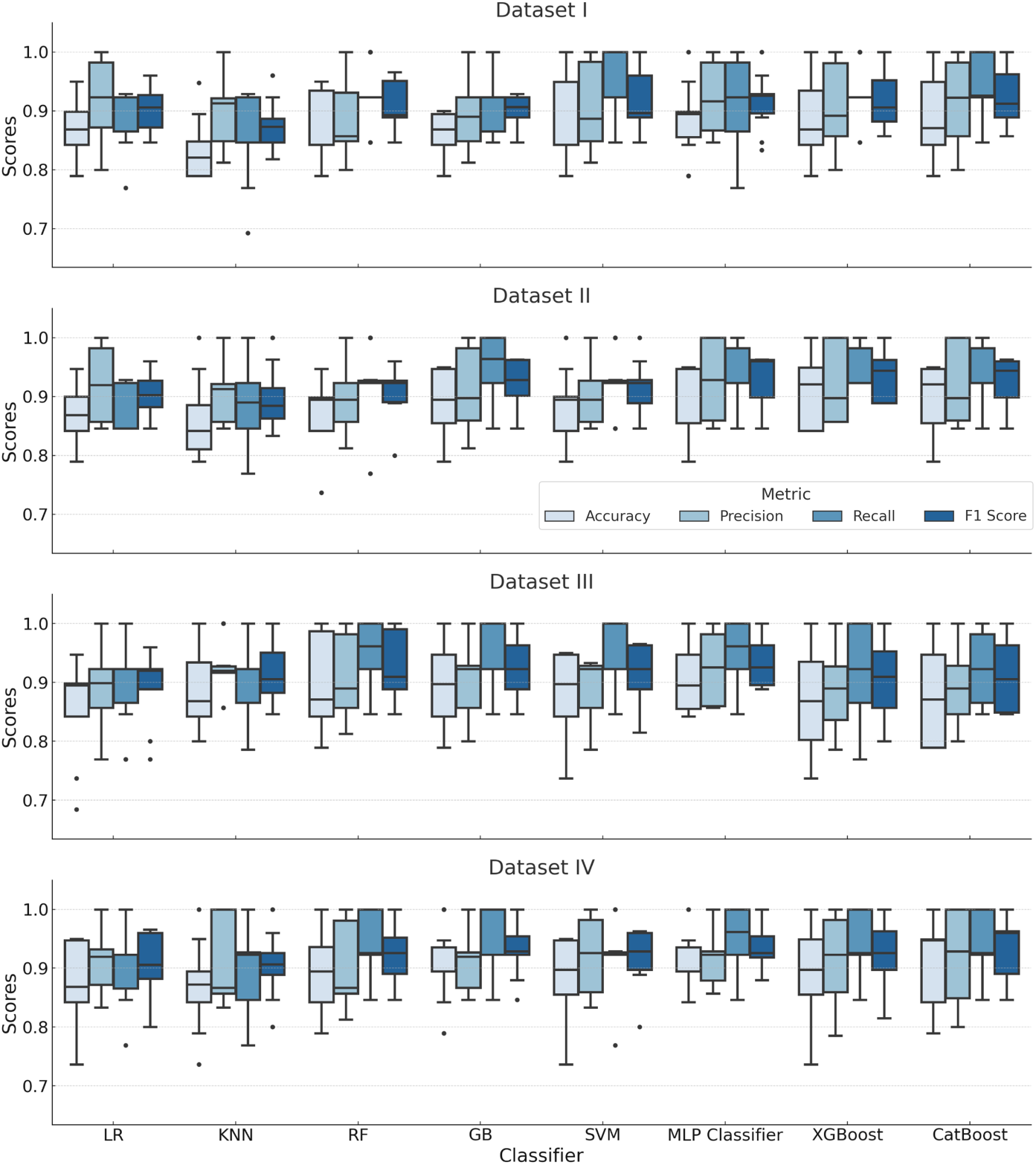
Comparison of F1 score performance across all ML algorithms.

#### 3.2.2. Multi-temporal classification

We used imagery from 2023 to determine the most effective season for distinguishing between the two species. In addition to the full dataset that contains all the spectral band and index values from all months (Table 3, Dataset I), we added three datasets. The results from feature importance testing indicated that 76 features were important for species discrimination. Thus, one dataset contained all 76 features (Table 3, Dataset II). As some of the months had none or few important features, we decided to create two additional datasets from the months with the highest number of informative features. Thus, one of the datasets contained data from April and November, and the other data from April, September, October and November, using only features which significantly distinguished the species across those months (Table 3, dataset III and IV, respectively).

**Table 3.** Description of the datasets used in the study.

| Dataset | Description | Number of Bands | Number of Indices |
| --- | --- | --- | --- |
| Dataset I | Full dataset | 72 | 135 |
| Dataset II | Feature selection | 23 | 53 |
| Dataset III | April and November | 7 | 22 |
| Dataset IV | April, September, October, and November | 11 | 38 |

Each dataset was used independently for training and testing to ensure accurate performance evaluation. All of the datasets gave F1 scores > 0.90 (Table 4). The differences between the datasets were minor, indicating that this approach also yields good results from smaller datasets, such as Datasets III and IV (Figure 4). Dataset I consistently showed higher performance across all classifiers, while Dataset II performed slightly worse but still maintained good performance, indicating that the statistically different features captured most of the essential information. Dataset I, with 207 features, has higher dimensionality, providing more information for the classifiers and helping to achieve slightly higher and more consistent classification performance, but perhaps at a greater risk of overfitting to this specific dataset. Dataset II outperformed Datasets III and IV, consistent with expectations, given that Dataset II contained significantly different features and was designed to retain the most informative features for the classification of the two species. The combination of spectral bands and spectral indices also showed a slight improvement in the results, improving the performance by five per cent in some of the ML algorithms (SVM, MLP, see section 7, results-datasets.xlxs).

**Table 4.** Performance metrics for different models across different datasets, mean values from the 10-fold cross-validation.

| Model | Dataset I |  |  |  | Dataset II |  |  |  | Dataset III |  |  |  | Dataset IV |  |  |  |
| --- | --- | --- | --- | --- | --- | --- | --- | --- | --- | --- | --- | --- | --- | --- | --- | --- |
|  | Acr | Prec | Rec | F1 | Acr | Prec | Rec | F1 | Acr | Prec | Rec | F1 | Acr | Prec | Rec | F1 |
| LR | 0.869 | **0.919** | 0.893 | 0.904 | 0.869 | 0.917 | 0.893 | 0.904 | 0.854 | 0.896 | 0.892 | 0.893 | 0.879 | 0.917 | 0.908 | 0.911 |
| KNN | 0.833 | 0.903 | 0.854 | 0.874 | 0.859 | 0.901 | 0.893 | 0.896 | 0.886 | **0.924** | 0.909 | 0.915 | 0.869 | 0.913 | 0.901 | 0.904 |
| RF | 0.869 | 0.889 | 0.931 | 0.908 | 0.869 | 0.892 | 0.924 | 0.906 | 0.901 | 0.908 | **0.954** | 0.930 | 0.890 | 0.903 | 0.947 | 0.922 |
| GB | 0.859 | 0.892 | 0.908 | 0.898 | 0.895 | 0.911 | **0.947** | 0.925 | 0.895 | 0.906 | 0.946 | 0.925 | **0.906** | 0.914 | **0.954** | **0.932** |
| SVM | 0.885 | 0.904 | 0.938 | 0.918 | 0.880 | 0.905 | 0.924 | 0.913 | 0.880 | 0.891 | 0.938 | 0.914 | 0.890 | 0.919 | 0.924 | 0.920 |
| MLP | 0.885 | 0.920 | 0.916 | 0.915 | 0.901 | **0.927** | 0.931 | 0.928 | 0.911 | 0.922 | **0.954** | **0.936** | **0.906** | **0.921** | 0.946 | 0.931 |
| Xgb | 0.879 | 0.902 | 0.931 | 0.914 | **0.906** | 0.922 | **0.947** | 0.933 | 0.864 | 0.884 | 0.923 | 0.902 | 0.896 | 0.913 | 0.939 | 0.925 |
| CatB | **0.896** | 0.910 | **0.947** | **0.926** | 0.901 | 0.922 | 0.939 | **0.929** | 0.874 | 0.898 | 0.924 | 0.910 | 0.901 | 0.917 | 0.947 | 0.930 |

To test for reliable performance, we conducted a controlled manipulation of the data by randomly altering the class labels. This and the following additional analyses were conducted using Dataset I. This led to a marked decrease in performance, yielding a significantly lower F1-score of 0.28 - 0.49. (Table S4; see section 7, supplementary and additional-results.xlsx). Given that our dataset contains about twice as many European as Caucasian beech data points, we furthermore equalized the class distribution by removing a portion of the European beech pixels (see section 2.3.3). While some algorithms maintained consistent performance, others showed a slight decrease when trained on the more balanced but reduced dataset (Table S5). Specifically, the Random Forest, Support Vector Machine, and MLP classifiers maintained similar performance as for the full dataset (for example, Random Forest: full dataset 0.87, balanced dataset: 0.86). However, the other metrics were lower with the balanced data (example, Random Forest: full dataset F1 score 0.91, balanced dataset F1 score: 0.86) (see section 7, additional-results.xlsx). Overall, F1 scores on this balanced dataset ranged from 0.78 to 0.87, versus 0.87-0.93 on the original Dataset I (Table 4).

We also compared our results from a UAV orthophoto taken on 14^th^ July 2022 over Allenwiller to the 18^th^ July 2022 results from the PlanetScope SuperDove data. The UAV data has one extra spectral band and higher spatial resolution. For the PlanetScope classification, the 8 spectral bands were used. We conducted the classification using a single pixel per tree for the sake of comparison. This resulted in an F1 score ranging from 0.56 - 0.72 for the PlanetScope image, and 0.71 - 0.81 for the UAV orthophoto (Figure 5C; Table S6, see section 7, additional-results.xlsx).

**Figure 5:**
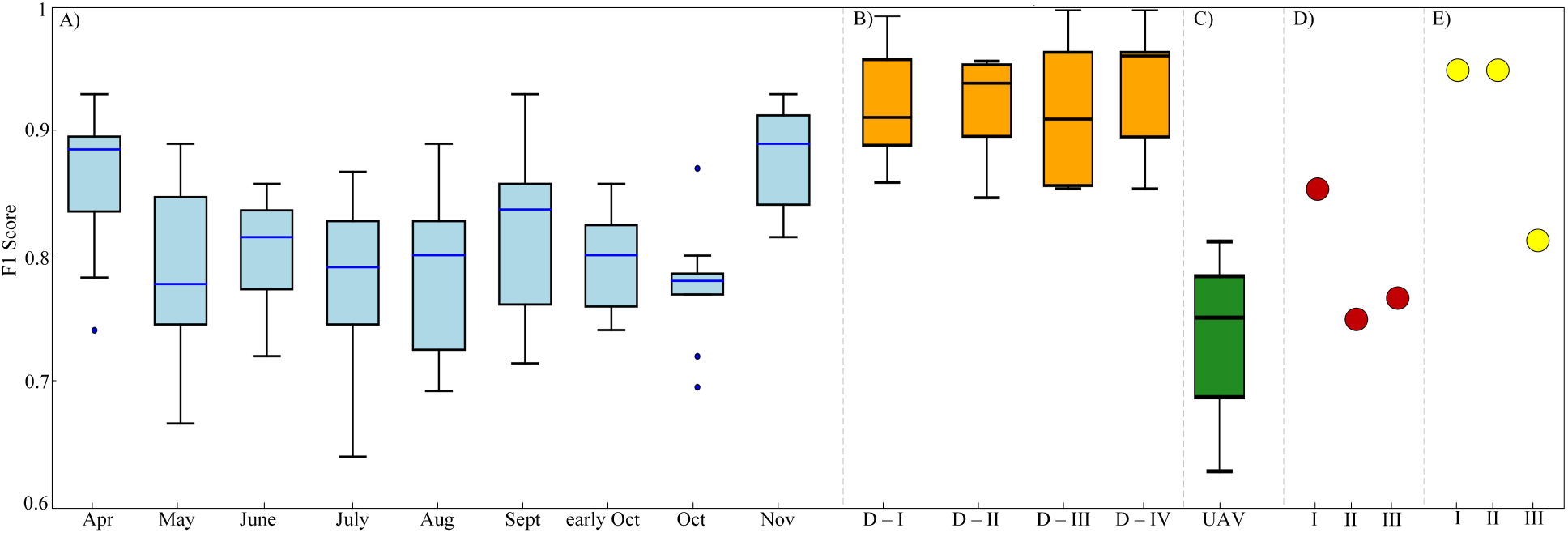
Comparison of classification performance (F1 scores) using different data. For A-C, results of the CatBoost model are shown as boxplots and for E, results of MLP are shown. Results from all models are given in Tables 5 and 6. A) Single date classification (April - November); B) Multi-temporal classification (D stands for Dataset, Dataset I - IV); C) UAV image results. D) Cross-temporal validation (I - 2021 training, 2022 testing; II - 2021 training, 2023 testing; III - 2023 training, 2022 testing.) E) Real-world application results over Wäldi (I - 2021; II - 2022; III - 2023).

**Table 5.**
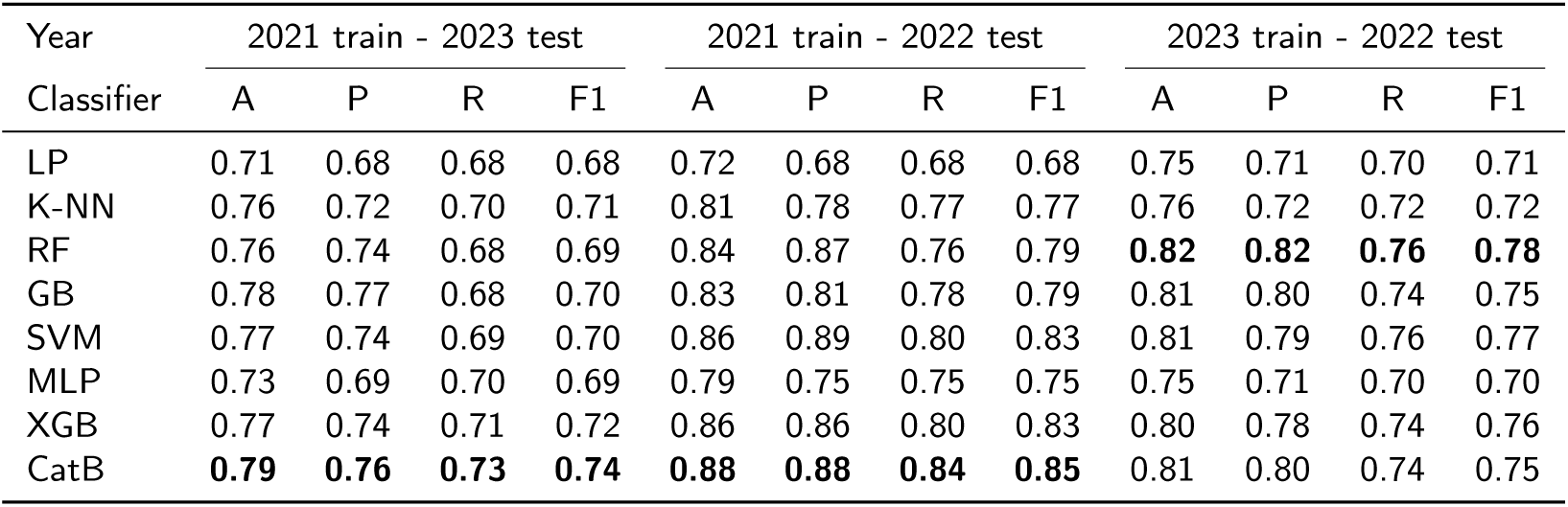
Comparative performance metrics (Accuracy - A; Precision - P; Recall - R; F1 - F1 Score) of different classifiers across three years.

| Year | 2021 train - 2023 test |  |  |  | 2021 train - 2022 test |  |  |  | 2023 train - 2022 test |  |  |  |
| --- | --- | --- | --- | --- | --- | --- | --- | --- | --- | --- | --- | --- |
| Classifier | A | P | R | F1 | A | P | R | F1 | A | P | R | F1 |
| LP | 0.71 | 0.68 | 0.68 | 0.68 | 0.72 | 0.68 | 0.68 | 0.68 | 0.75 | 0.71 | 0.70 | 0.71 |
| K-NN | 0.76 | 0.72 | 0.70 | 0.71 | 0.81 | 0.78 | 0.77 | 0.77 | 0.76 | 0.72 | 0.72 | 0.72 |
| RF | 0.76 | 0.74 | 0.68 | 0.69 | 0.84 | 0.87 | 0.76 | 0.79 | **0.82** | **0.82** | **0.76** | **0.78** |
| GB | 0.78 | 0.77 | 0.68 | 0.70 | 0.83 | 0.81 | 0.78 | 0.79 | 0.81 | 0.80 | 0.74 | 0.75 |
| SVM | 0.77 | 0.74 | 0.69 | 0.70 | 0.86 | 0.89 | 0.80 | 0.83 | 0.81 | 0.79 | 0.76 | 0.77 |
| MLP | 0.73 | 0.69 | 0.70 | 0.69 | 0.79 | 0.75 | 0.75 | 0.75 | 0.75 | 0.71 | 0.70 | 0.70 |
| XGB | 0.77 | 0.74 | 0.71 | 0.72 | 0.86 | 0.86 | 0.80 | 0.83 | 0.80 | 0.78 | 0.74 | 0.76 |
| CatB | **0.79** | **0.76** | **0.73** | **0.74** | **0.88** | **0.88** | **0.84** | **0.85** | 0.81 | 0.80 | 0.74 | 0.75 |

**Table 6.**
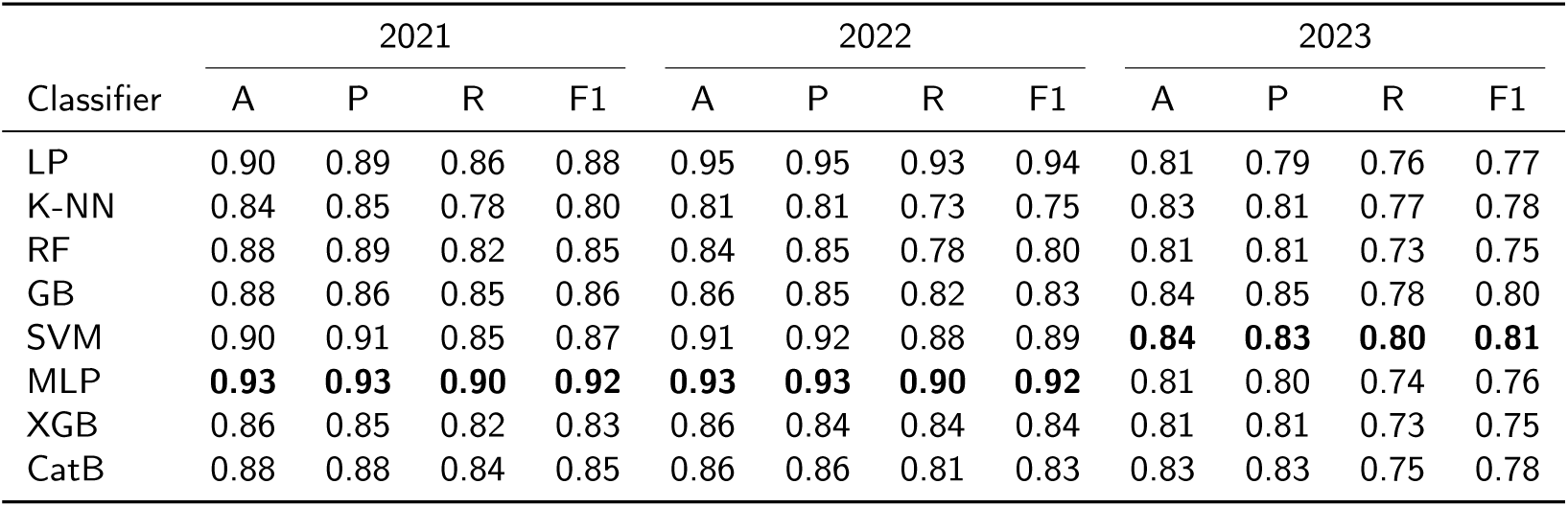
Performance metrics (Accuracy - A; Precision - P; Recall - R; F1 - F1 Score) of different classifiers over three years over the Wäldi study area.

| Classifier | 2021 |  |  |  | 2022 |  |  |  | 2023 |  |  |  |
| --- | --- | --- | --- | --- | --- | --- | --- | --- | --- | --- | --- | --- |
|  | A | P | R | F1 | A | P | R | F1 | A | P | R | F1 |
| LP | 0.90 | 0.89 | 0.86 | 0.88 | 0.95 | 0.95 | 0.93 | 0.94 | 0.81 | 0.79 | 0.76 | 0.77 |
| K-NN | 0.84 | 0.85 | 0.78 | 0.80 | 0.81 | 0.81 | 0.73 | 0.75 | 0.83 | 0.81 | 0.77 | 0.78 |
| RF | 0.88 | 0.89 | 0.82 | 0.85 | 0.84 | 0.85 | 0.78 | 0.80 | 0.81 | 0.81 | 0.73 | 0.75 |
| GB | 0.88 | 0.86 | 0.85 | 0.86 | 0.86 | 0.85 | 0.82 | 0.83 | 0.84 | 0.85 | 0.78 | 0.80 |
| SVM | 0.90 | 0.91 | 0.85 | 0.87 | 0.91 | 0.92 | 0.88 | 0.89 | **0.84** | **0.83** | **0.80** | **0.81** |
| MLP | **0.93** | **0.93** | **0.90** | **0.92** | **0.93** | **0.93** | **0.90** | **0.92** | 0.81 | 0.80 | 0.74 | 0.76 |
| XGB | 0.86 | 0.85 | 0.82 | 0.83 | 0.86 | 0.84 | 0.84 | 0.84 | 0.81 | 0.81 | 0.73 | 0.75 |
| CatB | 0.88 | 0.88 | 0.84 | 0.85 | 0.86 | 0.86 | 0.81 | 0.83 | 0.83 | 0.83 | 0.75 | 0.78 |

##### Cross-temporal validation

We then employed a cross-temporal validation strategy in which we used one year of satellite imagery for training and a dataset from another year for testing. For comparison, we used data from the same year for both training and testing, for which we obtained an F1 score of 0.86 for K-NN and > 0.93 for the other ML classifiers (see section 7, additional-results.xlsx). We trained our models using data from 2021 and tested them with data from 2022 and 2023. We then reversed this process, training on data from 2023 and testing on data from 2022. This approach allowed us to evaluate performance across different time periods and the possibility of over-fitting to a specific year’s data. Training with 2021 data and testing with data from 2022 produced F1 scores ranging from 0.68 - 0.74, whereas training on 2021 data and testing on 2023 data produced F1 scores ranging from 0.78 - 0.85; similarly, training on 2023 data and testing on 2022 data yielded F1 scores ranging from 0.70 - 0.78 (Table 5; Figure 5D), see section 7, additional-results.xlsx).

#### 3.2.3. Real-world application

Training on the Allenwiller dataset and testing on the Wäldi dataset performed well for 2021 and 2022 across all machine learning classifiers, with F1 scores ranging from 0.80 - 0.92 for 2021, and 0.75 - 0.92 for 2022 (Table 6). Performance for 2023 was lower, but maintained F1 values between 0.75 and 0.81 (Table 6, Figure 5E).

As expected, performance was markedly lower when we trained models on the smaller Wäldi dataset and tested them on the larger Allenwiller dataset. In 2021, the performances were highly variable, with F1 scores ranging from 0.39 to 0.91. Random Forest, SVM, and XGBoost consistently yielded the highest scores, each having an F1 score of 0.81. In 2022, similar levels of performance were only maintained with SVM (F1 = 0.81), while other methods showed significantly lower F1 scores of 0.45-0.69. For 2023, although scores were generally higher (0.81-0.87), they were uniformly distributed across the methods. In addition, some models yielded a “perfect” accuracy and F1 score of 1 in 2022 (MLP) and 2023 (LR, CatB), which raises concerns about overfitting or class imbalance or errors in model evaluation (Table S7).

Moreover, we employed the pixel prediction approach on a part of the Wäldi study area, avoiding reliance on individual tree data. We used only satellite imagery as input, resulting in an image with two classes: European and Caucasian beech. We specifically selected areas known to contain samples from both species. The results are illustrated in Figure 6A.

**Figure 6:**
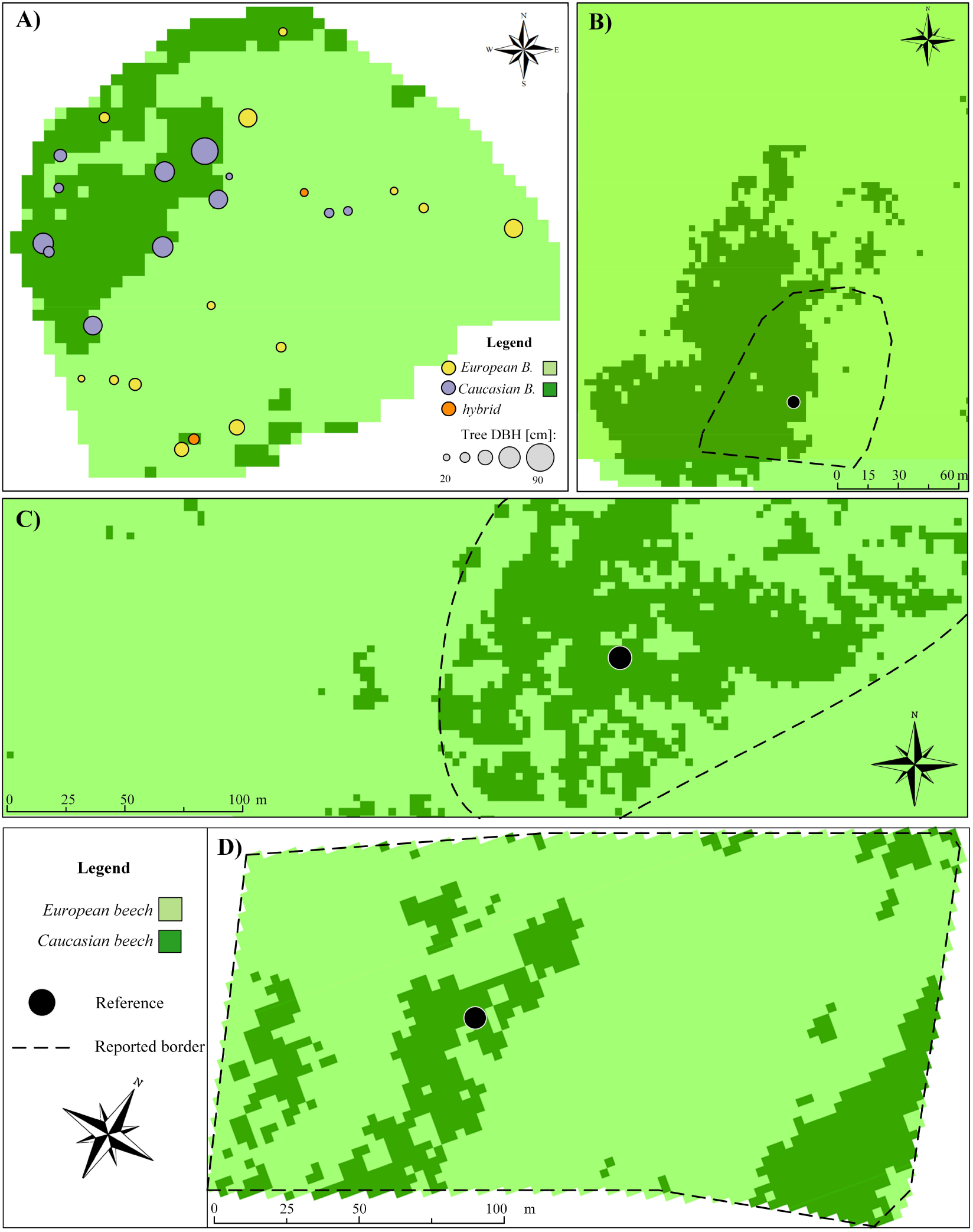
Pixel prediction for A) Wäldi; B) Leiselheim; C) Kirkel; D) Vorsenz.

Using additional data from three sites, our prediction results indicate the presence of Caucasian beech in all three study areas (Figure 6B-D), close to the areas indicated by foresters. The results from Leiselheim showed presence in the probable area, with a slight shift to the west. The results from Kirkel fitted directly with the probable area. It should be noted that the probable area continues north-east. However, we expect our partial prediction is owing to cloud cover over some of the images in 2023. Vorsenz is the largest sampled area in the previous study, where, according to the authors, there is a large number of individual Caucasian beech trees within the defined stand.

## 4. Discussion

Our study demonstrates that it is feasible to distinguish the closely related, hybridizing, and optically similar European and Caucasian beech using satellite data, allowing to make predictions rapidly across different forest stands. Spectral analysis conducted with 8-band PlanetScope satellite imagery across multiple years revealed that these species could best be distinguished in the spring and fall by leveraging phenological differences at sites where they co-occur. A previous study reported the SWIR domain to be the most informative region for discriminating Caucasian and European beech, but this was determined from single-timepoint, mid-season samples (July, (D’Odorico et al., 2023)). However, this study aligns with findings from macrophenology research, which highlights the importance of leveraging broad-scale, multi-temporal data, as phenological responses can very significantly with environmental cues such as temperature and photo period (Gallinat, Ellwood, Heberling, MillerRushing, Pearse, and Primack, 2021). Our study thus informs monitoring strategies where SWIR bands are not available, which is usually the case when using multispectral sensors. Moreover, our results are supported by the multiscale approach demonstrated in the study by Berra, Gaulton, and Barr (2019), where ground-based observations, UAV-data, and satellite imagery were compared to assess spring phenology in temperate woodlands. Our results furthermore indicate that the spectral signatures from 8-band imagery in July actually provide the poorest distinction between European and Caucasian beech. This was also confirmed with the feature selection analysis, where none of the spectral band or index values from July were found to differ significantly. Instead, our results strongly support the utility of multispectral, multi-temporal data that is readily available from satellite sensors to capture phenological differences that can distinguish similar tree species or possibly even provenances that trait divergence in bud break (Aitken and Bemmels, 2016; Gallinat et al., 2021).

### 4.1. Phenology and classification accuracy

Single-date classification tests revealed that April and November were the most informative months for distinguishing Caucasian from European beech in our study area. These months likely correspond to key phenological shifts that are captured in spectral signatures, thus aiding in the discrimination process (Figure 3, (Siachalou, Mallinis, and Tsakiri-Strati, 2015)). In contrast, models trained and tested on data from June, July and August performed worse. This suggests that the spectral signatures during these months are less distinct between the species, likely because differences in their canopy optical properties under leaf-on conditions mid-season are small compared to the difference in optical properties between leaf-on and leaf-off conditions that can be captured in spring and fall as a result of differences in their leafing-out phenologies. This variability in classification accuracy across months reflects broader patterns identified in macrophenology, where phenological shifts driven by chilling, growing degree days, and photoperiod can lead to significant interannual variability in species responses (Gallinat et al., 2021). The importance of selecting optimal times for data acquisition is further highlighted by Berra et al. (2019), who demonstrated how phonological stages captured through different observational scales (ground, UAV, and satellite) can significantly influence classification accuracy. In the intervening period during which both species had primarily mature leaves, higher-dimensional data, such as leaf contact measures with a higher-resolution spectrometer, may help to better distinguish them (D’Odorico et al., 2023). Kurz et al. (2023) reported earlier spring phenology of adult Caucasian beech trees at sites where Caucasian beech was planted into European beech stands. We were not able to capture continuous spring phenological differences due to limited cloud-free PlanetScope imagery. However, during the autumn months, European beech exhibited significantly higher reflectance values across most spectral bands than Caucasian beech at one of these sites, corresponding to the apparent earlier leaf abscission of Caucasian beech, perhaps resulting from an earlier bud break in spring. Thus, late autumn observations were the most informative in our main training and testing dataset. The variability in classifier performance across months in our study underscores the necessity of selecting optimal times for data acquisition, which can significantly impact the performance and reliability of species classification. While the timing of image capture is known to be critical for vegetation classification tasks (Dudley, Dennison, Roth, Roberts, and Coates, 2015; Zhong, Hu, and Zhou, 2019), we highlight the importance of phenological differences in particular for distinguishing these optically similar species.

Yet phenology may not always be informative for distinguishing highly similar, co-occurring species, and not even for distinguishing Caucasian and European beech in all circumstances. Plant phenology responds to environmental conditions and its importance in distinguishing species will depend on the plasticity of their phenology and the range of environmental conditions included (Wolkovich, Chamberlain, Buonaiuto, Ettinger, and Morales-Castilla, 2022; Buonaiuto and Wolkovich, 2021). Leaf phenology of beech and other tree species is known to be influenced by chilling requirements, growing degree days, as well as the photoperiod, which can differ by several weeks across environmental gradients, as well as from year to year (see e.g. (Lukasová, Vido, Škvareninová, Bičárová, Hlavatá, Borsányi, and Škvarenina, 2020; Müller, Kempen, Finkeldey, and Gailing, 2020; Walde, Wu, Fox, Baumgarten, Fu, Wang, and Vitasse, 2022)). Along these lines, cross-temporal validation revealed that models trained on 2021 data and tested on data from 2022 and 2023 performed consistently well, indicating that the approach effectively captured stable spectral features across years. However, a noticeable decline in performance occurred when models trained on 2023 data were tested against the 2022 dataset. This decrease may be due to several factors, including phenological or environmental variations not captured in the 2023 training data but present in the 2022 data.

Our main training and testing site for this study, Allenwiller, happens to be at an intermediate latitude among our various test sites (see Figure 1). While Allenwiller was chosen because it hosted the largest number of genotyped and geolocated trees, it may also be that the phenology at Allenwiller is sufficiently representative of all sites used, whereas the phenology at Wäldi (southernmost) or Kirkel (northernmost) may be least representative. It is possible that models trained for trees under different temperature regimes, e.g. across latitudes or across years (especially given climate change), would identify other time points with the best classification power, or other features. Furthermore, it is unclear whether the phenological differences observed between species under shared environmental conditions as in our study (located in the same forest stands) will be helpful to distinguish the species across larger environmental gradients, for example when adding sites across Europe or attempting predictions across the boundaries of the species’ distributions and their natural hybrid zone. We therefore recommend that future work in this direction incorporate models of the species’ phenologies dependent on relevant environmental factors to guide the selection of imagery used for classification as well as feature selection. In the absence of such predictive information about phenology across the study species, area and timeframe, we currently recommend to use SVM, which was both the most robust of the ML models to variation in class-balance and also showed the best temporal generalizability.

In any case, multi-temporal data is still likely to identify the overall most distinguishing features across growing seasons. Our findings emphasize the importance of leveraging multi-temporal data to enhance classification accuracy, especially for vegetation classification tasks where canopy optical properties are not strongly distinguishing. Furthermore, other studies have indicated that fall phenology is useful even for distinguishing more optically distinct species, including separating evergreen from deciduous species, but also distinguishing among deciduous species in mixed forests (Vitasse, Bresson, Kremer, Michalet, and Delzon, 2010).

### 4.2. Spatial resolution and classification accuracy

Further analysis involved comparing the July results from the PlanetScope SuperDove data (8 bands), taken on July 18, 2022, with a UAV orthophoto taken on July 14, 2022. The UAV data, which includes extra spectral bands and higher spatial resolution, provided a valuable comparison point (see section 7, additional-results.xlsx). However, the classification performance was similar, indicating that the additional spectral and spatial resolution did not substantially improve classification performance for this task. Our results align with previous findings, where despite the higher spatial resolution of UAV images, the performance of satellite images was still competitive (Ruwaimana, Satyanarayana, Otero, M. Muslim, Syafiq A, Ibrahim, Raymaekers, Koedam, and Dahdouh-Guebas, 2018). Given the effort required to generate UAV data, satellite data may often be preferable. It should be noted that the UAV footage was acquired outside of the most informative time points and was multispectral, i.e., had lower spectral resolution than the leaf contact measurements taken by Kurz et al. (2023), which limited its ability to detect subtle physiological differences. To maximize the information such imagery can provide about the species, we suggest that UAV footage be collected during key phenological transitions as determined by satellite imagery, such as early spring and/or late fall, when physiological changes are more pronounced and detectable.

### 4.3. Representativeness and information content of input data

Datasets incorporating data from multiple months, particularly those with data from every month (Dataset I) or from April, September, October, and November (Dataset II), generally demonstrated the highest classification performance. The inclusion of spectral indices in these multi-temporal datasets improved performance by up to 5%, perhaps due to having greater sensitivity to detect subtle differences because the indices are normalized (Thapa, García-Millán, and Eklundh, 2021; Praticò, Solano, Fazio, and Modica, 2021; Chrysafis, Korakis, Kyriazopoulos, and Mallinis, 2020). Despite the smaller number of features in Datasets III and IV, high F1 scores over 0.90 were achieved, indicating that feature selection was effective. We therefore recommend feature selection in order to avoid overfitting (see also subsection 4.4, below).

To validate our results and ensure robustness, we tested the model by randomly altering class labels, which, as expected, caused a significant drop in performance. The drop in performance with artificially mixed data supports that our high performance rates are not due to chance (Japkowicz and Stephen, 2002; Kittlein, Mora, Mapelli, Austrich, and Gaggiotti, 2021; Chawla, Bowyer, Hall, and Kegelmeyer, 2002). Additionally, addressing dataset imbalance by removing some European beech data led to varied results across algorithms, with some maintaining performance and others decreasing. Performance drops were on the order of ten percent on average for the balanced versus full dataset.

### 4.4. Limitations and Outlook

The evaluation of our approach across the Wäldi and Allenwiller datasets highlights key aspects of remote sensing-based classification, particularly the challenges of model generalization across different scales and resolutions. While the approach demonstrated high performance when trained and tested on the same year and forest, or trained on a larger dataset and tested on a smaller dataset, performance was variable and often poor when the model was trained on a smaller dataset and applied to a larger and more complex dataset. We point this out because it is a situation quickly encountered when using machine learning in ecological research; yet, it is not surprising and highlights the well-known importance of sufficient training data when building a classifier. A minimum of a few hundred samples is commonly recommended (Atkinson and Treitz, 2012), i.e., around the size of the Allenwiller dataset. We also note that the number of samples available for this study in Allenwiller (192 genotyped trees) is on the same order of magnitude as the 207 features used as a maximum for models (Dataset I in this study), which is why we recommend using reduced datasets with only the significantly differing features (e.g. Datasets II-IV). While model performance was slightly worse for Dataset I versus Dataset II, the advantage of Dataset I may be due to the very common problem of over-fitting.

To further improve the robustness of the approach we describe here will require more validation data than we currently have available, which is also a common limitation of machine learning and deep learning approaches. However, we demonstrate plausible predictions at the scale of satellite imagery tiles. These predictions should be tested, and the approach should be improved using additional point genotype data from the closely related Eurasian beech species (e.g. (Pfenninger, Langan, Feldmeyer, Fussi, Hoffmann, Granado, Hetzer, Šeho, Mellert, and Hickler, 2023; Czyż, Schmid, Hueni, Eppinga, Schuman, Schneider, Guillén-Escribà, and Schaepman, 2023; Kurz et al., 2023)). This will entail application to larger geographic areas such as the gradient of species ranges for the Eurasian beeches and their described hybrid zones, while we have so far only applied this approach to beech trees co-occurring in forest stands. However, because predictions based on one site (Allenwiller in eastern France) seemed to perform well at other sites in Switzerland and Germany, we think that our approach provides a good basis for expanding to larger geographic areas, incorporating the considerations discussed above, in particular incorporating predictive knowledge of leaf phenology.

Spatial resolution may present a greater challenge to the broader application of this approach, keeping in mind that we here have used data at relatively high spatial resolution in comparison to other sources of satellite multispectral data (3 m as opposed to 10-20 m; see e.g. (Ustin and Middleton, 2021)). The pixel prediction approach that allows classification of areas solely based on satellite imagery, without reliance on precise tree location data, as tested over the Wäldi data, performed well in areas where trees of the same species were grouped (which was the case for most of our data), as opposed to interspersed single and smaller trees (see Figure 6). This likely reflects the influence of spatial resolution. The PlanetScope imagery used, while sufficient for grouped trees, may not have the spatial resolution necessary to accurately discern individual trees within canopies of mixed or closely interspersed species. Future work should investigate the use of spectral de-mixing to improve performance (Gašparović, Dobrinić, and Pilaš, 2023).

## 5. Conclusion

This study highlights the potential of high-resolution satellite data in forest management and conservation efforts involving similar and even interbreeding species. Several machine learning models performed well to distinguish Caucasian from European beech in forest stands where the two species co-occur, with SVM demonstrating the most robust performance across different scenarios. Our pixel-based approach does not rely on precise tree coordinates to generate predictions, which represents an advance, allowing predictions in areas where detailed ground data are unavailable. We have thereby generated predictions which should be validated in future ground-based studies.

The approach described here should be extended to the prediction of other species, both including a first step to distinguish Eurasian beech from non-beech species in mixed forest canopies, and then adding classification of the other three Eurasian beech species as required. The current approach is suitable for strongly beech-dominated forest canopies where Caucasian and European beech are expected to co-occur, which is the case for several beech forest stands in Europe where AGF is expected as a result of introductions in the past century (Kurz et al., 2023), and the aim for which this study was designed. However, the presence of sporadically occurring tree species that share similar phenological timings with beech, such as oak or chestnut, is likely to influence the performance of the approach as described here. Furthermore, where any other tree species are present, this approach will incorrectly classify them as either Caucasian or European beech. Expanding this approach to other tree species and ecological settings as discussed above would test its versatility and improve its utility for broader forest conservation and management efforts.

This study furthermore illustrates limitations imposed by the three-meter spatial resolution of the PlanetScope data used, which already represents high spatial resolution for available multispectral satellite data. While sufficient for analyzing large and grouped trees, this resolution likely does not capture the finer details required for smaller or sparsely distributed trees without incorporating methods to demix spectra, which should be investigated in future work. Deep learning-based methods to artificially improve spatial resolution by taking advantage of time series could also be considered (Fu, Liu, and Li, 2018), but may not be cost-effective when using commercial data sources such as PlanetScope and may also not be compatible with the need for phenological (temporal) resolution.

## Acknowledgements

This work was supported by a SwissForestLab grant (SFL-20 P6) to MCS and KC. KC is supported by an ERC Consolidator Grant (MyGardenOfTrees, 101003296). MCS acknowledges support from the URPP Global Change and Biodiversity program and the NOMIS Foundation project “Remotely Sensing Ecological Genomics”. GK acknowledges support by the CA19128 Cost Action. We thank Fabien Suter, Marie Mathys, Tis Voortman and Bernadett Virókné Fodor who conducted the field observations in Allenwiller, in November 2023. We thank the French National Forest Service (ONF), who manages the forest of Allenwiller, and the forest managers for Wäldi, Leiselheim, Kirkel, and Vorsenz that allowed the collection of ground-based data and provided additional information that made this study possible.

## 6. Data and code availability

All data and code necessary to replicate this work, and supplementary files are provided at https://zenodo.org/records/13283819 and https://github.com/ArianeMora/remseno. The coordinates of the single trees will be provided upon reasonable request.

## CRediT authorship contribution statement

**Gordana Kaplan:** Conceptualization, Data curation, Formal analysis, Investigation, Methodology, Resources, Software, Validation, Visualization, Writing - original draft, Writing - review & editing. **Ariane Mora:** Data curation, Formal analysis, Methodology, Resources, Software, Validation, Writing - review & editing. **Katalin Csilléry:** Conceptualization, Data curation, Funding acquisition, Investigation, Methodology, Project administration, Resources, Supervision, Writing - review & editing. **Meredith C Schuman:** Conceptualization, Data curation, Funding acquisition, Investigation, Methodology, Project administration, Resources, Supervision, Writing - review & editing.

## Notes

https://github.com/ArianeMora/remseno

https://zenodo.org/records/13283819

### Competing Interest Statement

The authors have declared no competing interest.

### Summary of Updates

some typos have been addressed in the introduction, discussion, and methods.

